# Unusual Ecofunctional Traits of *Endozoicomonas*: A Pan-Genomic Perspective

**DOI:** 10.1101/2025.03.06.641946

**Authors:** Sim Lin Lim, Ching-Hsiang Chin, Yu-Jing Chiou, Ming-Tsung Hsu, Pei-Wen Chiang, Hsing-Ju Chen, Yung-Chi Tu, Sen-Lin Tang

## Abstract

**Background:** *Endozoicomonas* is a widely distributed genus of marine bacteria, associated with various marine organisms, and recognized for its ecological importance in host health, nutrient cycling, and disease dynamics. Despite its significance, genomic features of *Endozoicomonas* remain poorly characterized due to limited availability of high-quality genome assemblies.

**Results:** In this study, we sequenced 5 novel *Endozoicomonas* strains and re-sequenced 1 known strain to improve genomic resolution. By integrating these 6 high-quality genomes with 31 others that were publicly available, we identified a distinct, coral-associated clade not recognized by the previous two-clade classification. Pan-genomic analysis revealed significant variation in genetic trait distribution among clades. Notably, *Endozoicomonas* lacks quorum sensing capabilities, suggesting resistance to quorum quenching mechanisms. It also lacks the ability to synthesize and transport vitamin B12, indicating that it is not a primary source of this nutrient for holobionts. A remarkable feature of *Endozoicomonas* is its abundance of giant proteins, ranging from 15 to 65 kbp. We identified 92 such proteins, which clustered into three major groups based on amino acid similarity, each associated with specialized functions, such as antimicrobial synthesis, exotoxin production, and cell adhesion. Additionally, we explored prophages and CRISPR-Cas systems. We found that *Endozoicomonas* acquired prophages from diverse sources via infection or other types of gene transfer. Notably, CRISPR-Cas sequences suggest independent evolutionary trajectories from both prophage acquisition and phylogenetic lineage, implying a potential influence of geographic or environmental pressures.

**Conclusions:** This study provides new insights into the genomic diversity of *Endozoicomonas* and its genetic adaptation to diverse hosts. Identification of novel genomic features, including deficiencies in B12 synthesis and quorum sensing, the presence of giant proteins, prophages, and CRISPR-Cas systems, underscores its ecological roles in various holobionts. These findings open new avenues for research on *Endozoicomonas* and its ecological interactions.

## Background

The genus *Endozoicomonas* comprises Gram-negative, rod-shaped marine bacteria that belong to the phylum *Pseudomonadota* (formerly *Proteobacteria*), belonging to the family *Endozoicomonadaceae* and the class *Gammaproteobacteria*. The discovery of the first species, *Endozoicomonas elysicola*, in a sea slug, *Elysia ornata*, in 2007, marked the beginning of extensive research into the ecological and functional roles of this genus [1]. Since then, *Endozoicomonas* has been identified in numerous marine organisms, including corals, sponges, soft corals, sea tunicates, fishes, and scallops [2–8]. It also exhibits remarkable adaptability, thriving in a broad range of marine environments, from deep-sea abyssal zones [9] to warm, photic regions [3].

Comprehensive 16S rRNA studies identify *Endozoicomonas* as a dominant member of various marine host-associated microbiomes, underscoring its ecological significance [1, 8, 10, 11]. Further 16S taxonomic profiling studies of coral associated microbiomes have demonstrated that *Endozoicomonas* strains not only dominate these communities [9, 12–15], but also display notable geographic specificity [16]. The discovery of complex interactions between corals and *Endozoicomonas* has sparked greater interest in marine microbiology. *Endozoicomonas* actively contributes to coral health and reef ecosystem stability. For example, declines in *Endozoicomonas* populations are frequently observed during coral bleaching, disease outbreaks, and other environmental stress events, coinciding with impaired coral health [17–19]. Conversely, corals recovering from bleaching events tend to harbor greater numbers of *Endozoicomonas*, suggesting roles in reef resilience and restoration [20].

Investigating the mutualistic functions of *Endozoicomonas* in marine organisms has proven challenging, largely due to the difficulty of cultivating and maintaining pure cultures under laboratory conditions [21]. *Endozoicomonas* isolates often require special and stringent cultivation conditions. For instance, *Endozoicomonas montiporae* is a slow-growing bacterium requiring specific sugar supplementation for growth [22]. Due to these limitations, researchers have turned to genomic approaches to gain insights into *Endozoicomonas* biology. Advances in sequencing technologies have facilitated assembly of *Endozoicomonas* genomes using sophisticated bioinformatics tools such as assemblers and metagenomic binning techniques [7, 23–25]. These genomic studies have shed light on the metabolic diversity and symbiotic interactions of *Endozoicomonas*, enabling a deeper understanding of its ecological roles.

Genomics and bioinformatics studies highlight the metabolic versatility and functional specialization of *Endozoicomonas* [10, 22, 26–30]. For example, these coral-dominant bacteria exhibit genetic traits that enable them to contribute to coral health by facilitating carbon, nitrogen, and sulfur acquisition, as well as degrading chitin and synthesizing secondary metabolites, including vitamins, cofactors, and hormones [10, 22, 26–30]. Experimental evidence has document key functions of three coral-dominant *Endozoicomonas*: *E. montiporae* strain CL-33^T^, *E. acroporae* and *E. ruthgatesiae* 8E. *Endozoicomonas montiporae* strain CL-33^T^ possesses the ability to degrade testosterone [22], while both *E. acroporae* and *E. ruthgatesiae* 8E metabolize dimethylsulfoniopropionate (DMSP) to dimethylsulfide (DMS) via the DddD CoA-transferase/lyase cleavage pathway, a process that helps corals mitigate thermal stress by scavenging free radicals [23, 29].

Comparative genomic studies of 7 *Endozoicomonas* genomes by Neave et al. (2017), 12 *Endozoicomonas* genomes by Alex and Antunes (2019), and 15 by Ide et al. (2022) laid the groundwork for understanding genomic diversity of *Endozoicomonas* [10, 26, 27]. These studies were not only limited by the *Endozoicomonas* genomes available at the time, but they were also constrained by limitations of next-generation sequencing (NGS) technologies, which primarily produced fragmented genome assemblies. This technical shortcoming restricted the ability to capture the full genomic diversity and features of the genus. The advent of third-generation sequencing (TGS) technologies has since revolutionized the field by enabling generation of high-quality, near-complete genome assemblies with vastly improved contiguity and accuracy. Unlike NGS, which struggles with repetitive regions and complex genomic architectures, TGS provides long-read sequencing that facilitates resolution of gene clusters, genomic structure/organization, gene orientation and order, regulatory regions, operons or prophages that were previously obscured. TGS technologies present an unprecedented opportunity for comprehensive comparative analyses, allowing researchers to uncover novel genomic and functional differences among strains and to unravel complex relationships between *Endozoicomonas* and its marine hosts.

In this study, we leveraged long-read HiFi sequencing technology to re-sequence 1 known strain for improved resolution and to assemble 5 high-quality genomes of novel *Endozoicomonas* strains isolated from stony and soft corals collected in Taiwan, the USA, and the Bahamas. By combining these new genomes with 31 publicly available *Endozoicomonas* genomic and metagenomic assemblies, we conducted the most comprehensive phylogenetic, pan-genome and genetic trait analyses of this genus to date. Additionally, we focused on novel genomic features that have been previously unexplored, such as giant proteins, prophages, and CRISPR-Cas systems.

## Materials and Methods

### Isolation, Sequencing, and Genome Assembly of 6 *Endozoicomonas* genomes

*Endozoicomonas.* sp KT9 was isolated from *Acropora* sp. (a scleractinian coral) in Kenting, Taiwan, *E.* sp 2B_B from *Acropora* sp. in the outer bay of Penghu main island, Taiwan, and *E.* sp PH6C from *Acropora muricata* in the inner bay of Penghu, Taiwan. All strains were isolated and cultivated following established protocols documented in the literature [31]. Stock cultures of *E. montiporae* CL-33^T^ from Southern Taiwan were obtained from a previous study [31], while *E. gorgoniicola* PS125 (Bimini, Bahamas) and *E. euniceicola* EF212 (Florida, USA) strains were sourced from the Westerdijk Fungal Biodiversity Institute (Utrecht, Netherlands) [7, 32].

For genome sequencing, high-quality genomic DNA was extracted using well-established methodologies [29]. Sequencing was performed on a PacBio Sequel IIe platform at Blossom Biotechnologies Inc. (Taipei, Taiwan), producing high-fidelity (HiFi) reads. Genome assemblies were generated using MetaFlye (v2.9.5) [33]. Genome assembly quality metrics, including genome completeness, contamination, and heterogeneity, were evaluated using CheckM [34]. We published genomic information *for E. gorgoniicola* PS125 and *E. euniceicola* EF212 very recently with only a brief, simple genomic analysis [7]. Both new genomes were reanalysed for detailed comparative genomic analyses in this study.

### Data Collection, 16S rRNA identification and Comparative Analysis of *Endozoicomonas* **Genomes**

A total of 73 publicly available *Endozoicomonas* genome assemblies and metagenome-assembled genomes (MAGs) were retrieved from the NCBI and RAST databases for comparative genomic analysis with the 6 genome assemblies generated in this study. Additionally, 3 bacterial genomes from different families, *Simiduia agarivorans* SA1 [35], *Marinobacterium aestuarii* ST58-10 [36], and *Cobetia marina* JCM21022 [37] were obtained from the NCBI database to serve as outgroup references. Detailed information regarding sources of these genome assemblies is presented in Supplementary Table S1. Full-length 16S rRNA sequences from the 82 genome assemblies were identified using Barrnap (https://github.com/tseemann/barrnap). Of these, 39 *Endozoicomonas* genome assemblies and 3 outgroup genome assemblies that contained at least 1 full-length 16S rRNA were retained for subsequent analyses. Meanwhile, 40 *Endozoicomonas* genome assembles that did not contain full-length 16S rRNA were discarded. For uniform gene annotation across all genome assemblies, Prokka was employed with default parameters to annotate the 42 genomes [38]. Genomic comparisons among *Endozoicomonas* genome assemblies were conducted using the OrthoANI tool to calculate Average Nucleotide Identity (ANI) percentages [39].

### Integrated Phylogenetic Analysis of 16S rRNA and Orthologous Genes

To construct a 16S rRNA phylogenetic tree, a single representative sequence per genome was selected for analysis. MUSCLE [40] was used for sequence alignment, and RAxML-NG [41] with the GTR+G substitution model was employed to construct a maximum-likelihood phylogenetic tree. Simultaneously, OrthoFinder [42] was utilized to identify single-copy orthologous genes across genome assemblies. Amino acid sequences of each ortholog were aligned using MUSCLE, and alignments were concatenated for subsequent phylogenetic analysis. A maximum-likelihood phylogenetic tree was generated using RAxML-NG with the LG+F+I+G4 substitution model. Both phylogenetic trees were visualized in MEGA11 to facilitate interpretation [43]. The initial analysis revealed that two *Endozoicomonas* genome assemblies, *Endozoicomonas* sp. G2_1 and *Endozoicomonas* sp. G2_2, were positioned as an outgroup with other outgroup reference genomes in both phylogenetic trees. As a result, both assemblies were excluded. Consequently, 37 *Endozoicomonas* genomes and 3 outgroup references were retained, and ANI-based genomic comparisons were revised.

### Pan-Genome Analysis and Genome-Wide Similarity of *Endozoicomonas*

Pan-genomic analysis of *Endozoicomonas* was performed using OrthoFinder to classify orthogroups among 40 genome assemblies [42]. Orthogroups were categorized into five groups: core (100% of genomes assemblies contain a given orthogroup), soft-core (95%– 99% of the genomes assemblies contain a given orthogroup), shell (15%–95% presence of the genomes assemblies contain a given orthogroup), cloud (2.5%–15% presence of the genomes assemblies contain a given orthogroup), and singletons. To examine relationships between orthogroups, hierarchical clustering based on orthogroup presence or absence (excluding singletons) was conducted in R [44]. Clustering results were arranged according to the phylogeny derived from single-copy core genes and visualized using PRISM. To assess genome-wide patterns and similarity among *Endozoicomonas* genome assemblies, non-metric multidimensional scaling (NMDS) was performed using the Vegan package in R [45].

### KEGG Module-Based Genetic Trait Analysis in *Endozoicomonas*

Protein sequences annotated with Prokka were processed using eggNOG-mapper to assign KEGG orthology (KO) terms to each genome assembly [46]. KO annotations enabled reconstruction of *Endozoicomonas* genetic traits by mapping onto KEGG modules. Genetic trait completeness was evaluated according to criteria outlined by Zoccarato et al. (2022): KEGG modules with three or more KO terms were deemed complete if up to one term was missing, whereas modules with fewer than three KO terms required all terms to be present [47]. R pheatmap was used to cluster and visualize genetic traits of *Endozoicomonas* with the “ward.D2” clustering method [44]. A dot plot was constructed to illustrate presence of genetic traits, aligned with the phylogeny of single-copy core genes. Furthermore, a radial plot was developed in Microsoft Excel to emphasize 13 specific interaction traits. Eleven interaction traits were derived from the genetic trait analysis, while the other 2 traits: Fe-siderophore synthesis and transport, were annotated with available KO terms due to the lack of a KEGG module (The trait was deemed complete in *Endozoicomonas* strains based on previously mentioned criteria). Detailed information regarding interaction traits available in these genome assemblies is presented in Supplementary Table S2.

### Analysis and 3D Modelling of *Endozoicomonas* Giant Proteins

Giant Proteins (GP) from *Endozoicomonas* genome assemblies were extracted based on Prokka annotations and allocated into 3 classes: small GP (>15 kbp), medium GP (>25 kbp) and large GP (>65 kbp). The Average Amino Acid Identity (AAI) between these proteins was calculated using EzAAI v1.2.3 [48], and pairwise AAI values were clustered with hierarchical clustering. Clustering results were visualized using PRISM. Secondary metabolite functions and protein domains were analyzed using InterProScan 5.70-102.0 [49]. The large GP domain associated with NRPS was further analyzed with antiSMASH 7.0 [50]. To predict three-dimensional structures of large GPs, the AlphaFold Server (version 2024.05.23) [51] was used. Each giant protein was divided into overlapping segments (700-residue overlap between consecutive segments) to meet the server’s amino acid size limitation of 2,800 residues. Resulting fragment structures were merged using PyMOL (v3.0.3) [52] to produce complete 3D model of the large GP.

### Prophage and CRISPR-Cas System Characterization in *Endozoicomonas*

Prophage regions in *Endozoicomonas* genome assemblies were identified using PHASTER [53]. Intact prophages underwent phylogenetic analysis with VIPtree [54], and their ANI percentages were calculated using the previously mentioned method. BLASTN [55] searches against the Non-Redundant (NR) database were used to characterize the gene content of intact prophages, identifying bacterial, eukaryotic, and phage-specific genes. Functional categorization of these genes was performed using COG annotations from eggNOG-mapper [46].

Identification of CRISPR-Cas systems was performed using CRISPRCasFinder [56]. Systems were considered complete if 1) the CRISPR array’s confidence level was 3 or 4; and 2) CRISPR arrays and *Cas* genes were within 4000 bp. For complete systems, consensus CRISPR sequences were aligned pairwise using EMBOSS needle [57] to compute sequence identity percentages. Hierarchical clustering of these sequence identities was executed in R pheatmap with default parameters [44], and results were visualized in PRISM.

## Results

### Six High-Quality *Endozoicomonas* Genome Assemblies

Five novel *Endozoicomonas* strains were isolated from diverse geographic regions and hosts. Two strains were obtained from soft corals (*Eunicea fusca* and *Plexaura sp.*) collected in Florida, USA, and Bimini, Bahamas [7, 32], whereas 3 strains were isolated from stony corals (*Acropora sp.*) in Taiwan. Additionally, the previously described *E. montiporae* CL-33^T^ was re-sequenced for comparative analysis. Genomic sequencing was performed using a PacBio Sequel IIe platform, followed by assembly with metaFlye [33]. This approach yielded high-quality genomes with 1–2 contigs per assembly, ranging in size from 6.34 to 7.48 Mbp (Table 1). Notably, resequencing improved the CL-33^T^ assembly from 3 contigs (5.43 Mbp) [22] to a single contig with a slightly larger genome size (5.44 Mbp). Quality evaluation using CheckM [34] demonstrated genome completeness of 98.71%–99.14% and low contamination levels (0.54%–2.19%), adhering to Genomic Standards Consortium guidelines for high-quality genome assemblies [58].

**Table 1.**
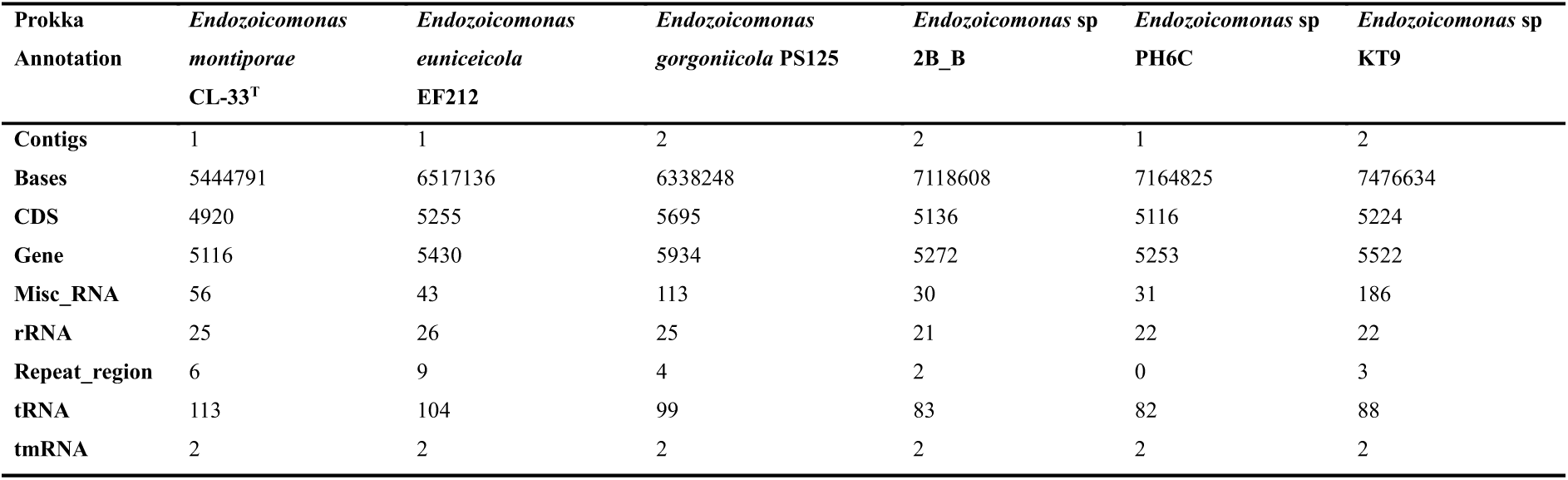
Genome assembly statistics of 6 *Endozoicomonas* genomes.

### Comparative Analysis of 37 Qualified *Endozoicomonas* Genomes

Comparative genomic analyses were conducted to contextualize the 6 *Endozoicomonas* genomes sequenced in this study in the broader diversity of *Endozoicomonas*. A dataset comprising 31 publicly available and qualified *Endozoicomonas* genomes (containing full-length 16S rRNA sequences) and 3 bacterial outgroup genomes (Supplementary Table S1) revealed that the newly sequenced genomes are well-polished, exhibiting markedly fewer contigs than existing assemblies, which typically range from 10 to over 1,000 contigs (Supplementary Figure S1A). The newly sequenced genomes were also significantly larger than previously reported *Endozoicomonas* genomes (Supplementary Figure S1B). However, gene content analyses showed no statistically significant differences in gene numbers (Supplementary Figure S1C). Across the entire dataset, a strong correlation was observed between genome size and gene number (R² = 0.92), suggesting conserved genomic scaling in this genus (Supplementary Figure S1D).

To investigate genomic relationships, average nucleotide identity (ANI) analysis was performed with OrthoANI [39] on the 6 newly sequenced genomes, 31 qualified *Endozoicomonas* genomes and 3 outgroups. ANI clustering (ANI > 80%) identified 3 major groups, with *E.* sp PH6C, *E.* sp 2B_B, and *E.* sp KT9 closely related to *E. ruthgatesiae* 8E, while *E. montiporae* CL-33^T^, *E. gorgoniicola* PS125, and *E. euniceicola* EF212 formed a distinct cluster (Figure 1). Notably, the grouping of *E. montiporae* CL-33^T^, *E. gorgoniicola* PS125, and *E. euniceicola* EF212 was independent of both geographic origin and host type, as CL-33^T^ was isolated from stony coral in Taiwan [22], whereas PS125 and EF212 originated from soft corals in North America [7].

**Figure 1.**
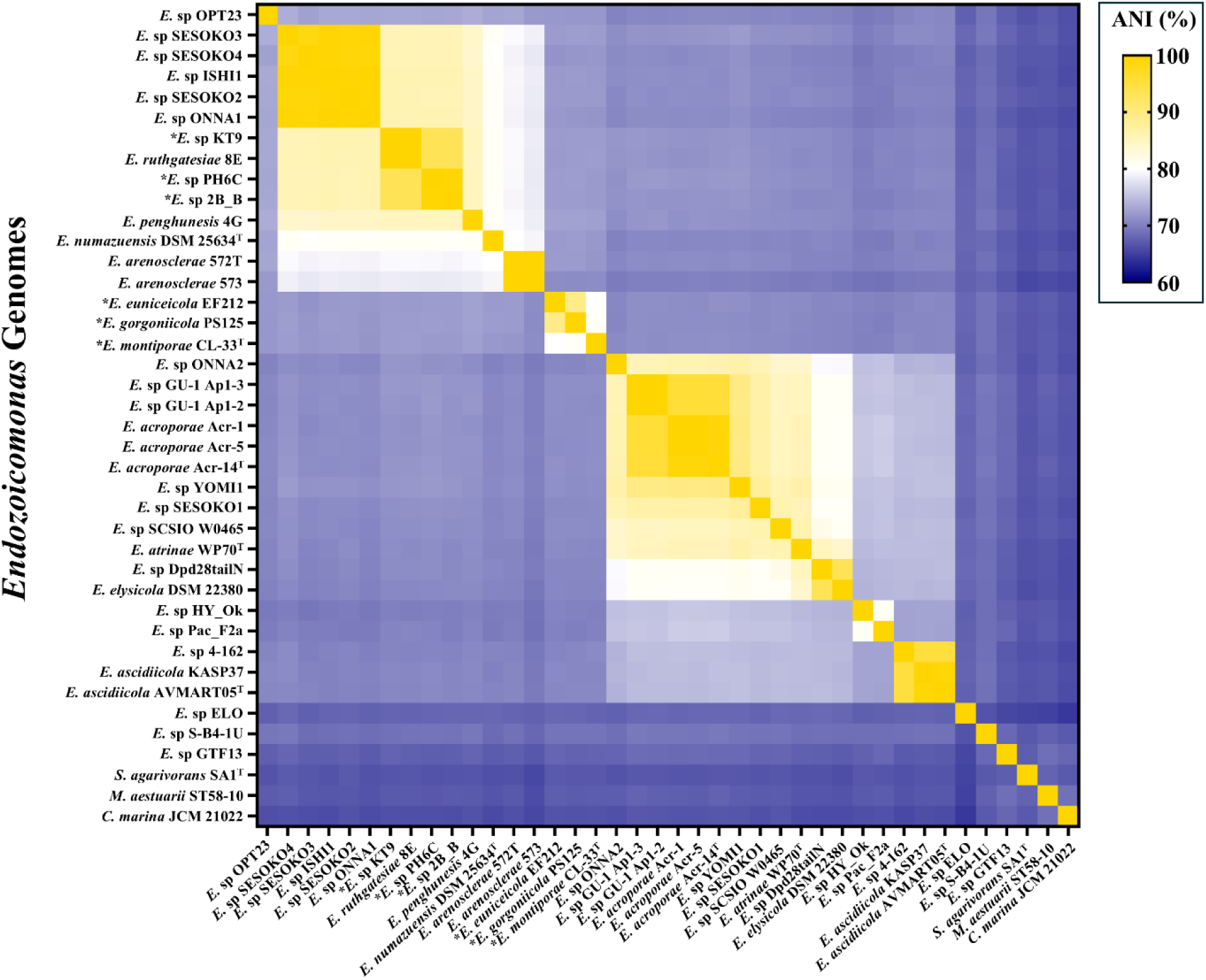
Average Nucleotide Identity (ANI) heatmap of 6 novel and re-sequenced *Endozoicomonas*, 31 publicly available *Endozoicomonas* genomes and 3 outgroups. *Endozoicomonas* strain positions were arranged according to single-copy core gene phylogeny. The * indicates *Endozoicomonas* strains sequenced in this study.

### Pan-Genome Analysis and Phylogenetic Insights into Genetic Relationships of 37 ***Endozoicomonas* Strains**

To evaluate whether the genomic structure inferred from ANI clustering aligns with phylogenetic relationships, we performed phylogenetic analyses using 2 major approaches. Phylogenies were constructed based on 16S ribosomal RNA (rRNA) gene sequences and a concatenated set of 57 bacterial core gene sets obtained from OrthoFinder output [42], supplemented by host information for 37 *Endozoicomonas* genomes (Figure 2). While both phylogenies identified 3 major clades, host associations within these clades differed slightly between the 2 methods. Bacterial core gene sets phylogeny grouped *Endozoicomonas* into 3 clades: the first (Clade 1) comprising strains associated with stony corals and sponges, the second (Clade 2) comprising strains associated with both soft and stony corals, and the third (Clade 3) comprising strains associated with multiple hosts excluding soft corals. In contrast, the 16S phylogeny yielded a first clade exclusively associated with stony corals, a second clade comprising strains associated with multiple hosts excluding soft corals, and a third clade associated with both soft and stony corals.

**Figure 2.**
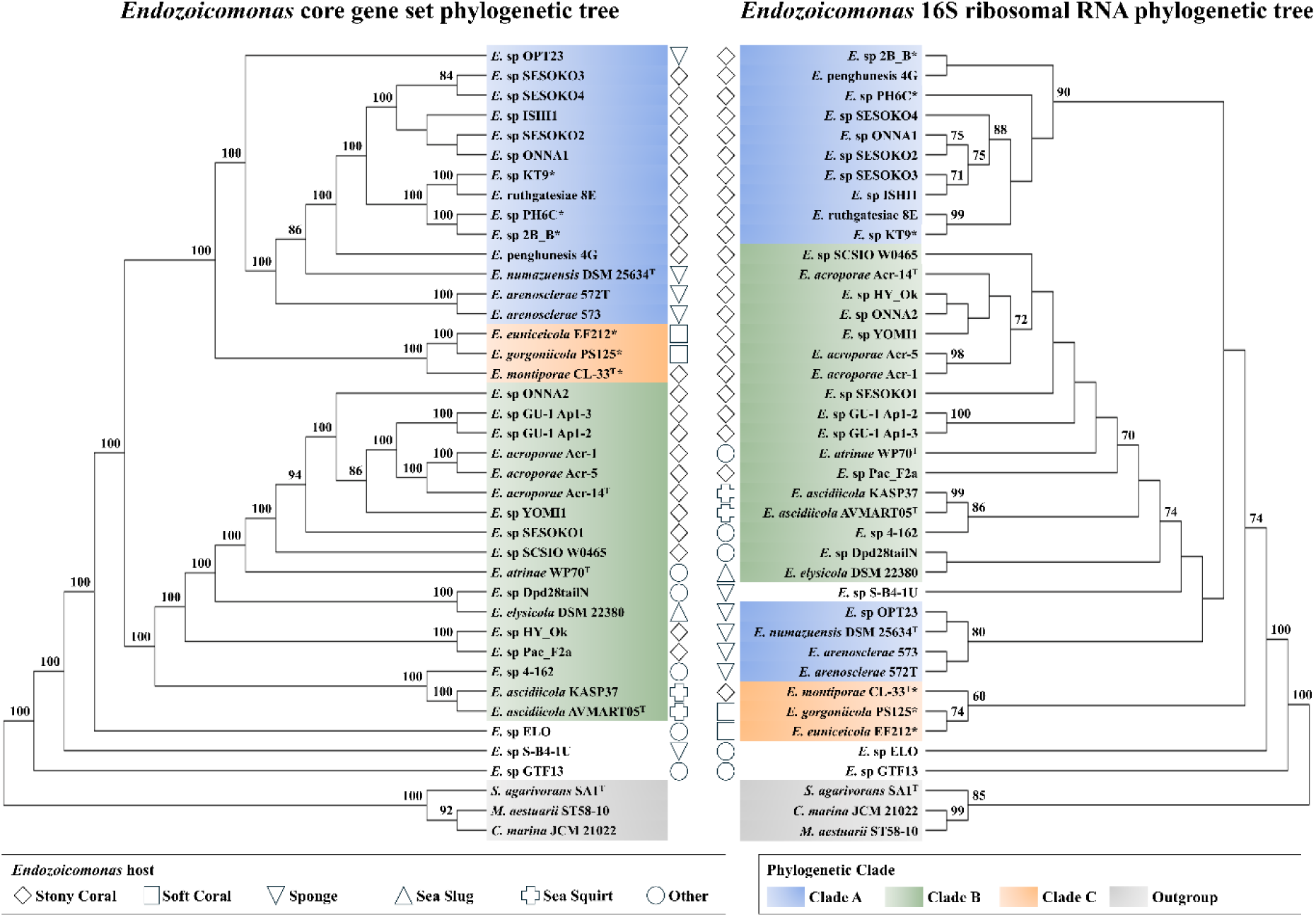
The *Endozoicomonas* phylogenetic tree constructed from single-copy core genes (left) and 16S ribosomal RNA (right). Both maximum-likelihood trees were calculated with RAxML-NG (GTR+G model). Blue, orange and green *Endozoicomonas* taxa represent three clades based on single-copy, core gene phylogeny. Grey taxa represent outgroups for *Endozoicomonas* genomes. * indicates *Endozoicomonas* strains sequenced in this study.

An orthogroup is a set of genes descended from a single gene in the last common ancestor of all species [42]. Therefore, it is useful for studying *Endozoicomonas* genomic diversity. This pan-genomic analysis was performed using Orthofinder [42]. Across the 37 *Endozoicomonas* genomes, 106 core orthogroups, 656 soft-core orthogroups, 4,838 shell orthogroups, 8,487 cloud orthogroups, and 11,069 singletons were identified. The orthogroup heatmap, mapped onto the single-copy, core gene phylogeny (Supplementary Figure S2A), revealed that the 3 clades exhibit distinct orthogroup compositions. Clade 1 and Clade 3 showed distinctive orthogroups, while Clade 2 appeared to share a mixture of orthogroups from Clade 1 and Clade 3, potentially explaining its intermediate phylogenetic position. Non-metric multidimensional scaling (NMDS) based on host factors further revealed that *Endozoicomonas* from stony corals could be divided into two subgroups, with one subgroup closely related to sponge-associated *Endozoicomonas*. Notably, *E. montiporae* CL-33^T^, *E. gorgoniicola* PS125, and *E. euniceicola* EF212 were positioned within a subgroup containing both stony and soft coral-associated bacterial strains, consistent with their intermediate genomic characteristics (Supplementary Figure S2B).

### Distinct Genetic Traits and Conserved Interaction Mechanisms Among *Endozoicomonas* **Clades**

Our analyses revealed that the 3 major clades of *Endozoicomonas* exhibit distinct gene content (Figure 2), suggesting substantial differences in their genetic traits and interaction mechanisms with both eukaryotic hosts and surrounding microbial communities. To further investigate these differences, we applied the genetic and trait identification framework of Zoccarato et al. 2022 [47], constructing a comprehensive genetic trait profile for 37 *Endozoicomonas* genomes and three non-*Endozoicomonas* outgroups based on KEGG modules (Figure 3A). A dot plot categorizing genetic traits identified four distinct groups: Category A traits, which occurred at low frequencies (<40%) in certain strains; Category B traits, predominantly associated with non-*Endozoicomonas* outgroups or distant *Endozoicomonas* strains (<10% frequency); Category C traits, shared by 20%–70% of *Endozoicomonas* strains; and Category D traits, core genetic features present in >80% of *Endozoicomonas* strains. Notably, the genetic trait distribution in Category A and Category C varied significantly among the 3 clades, providing further support for their differentiation as observed in the bacterial core gene set phylogeny.

**Figure 3.**
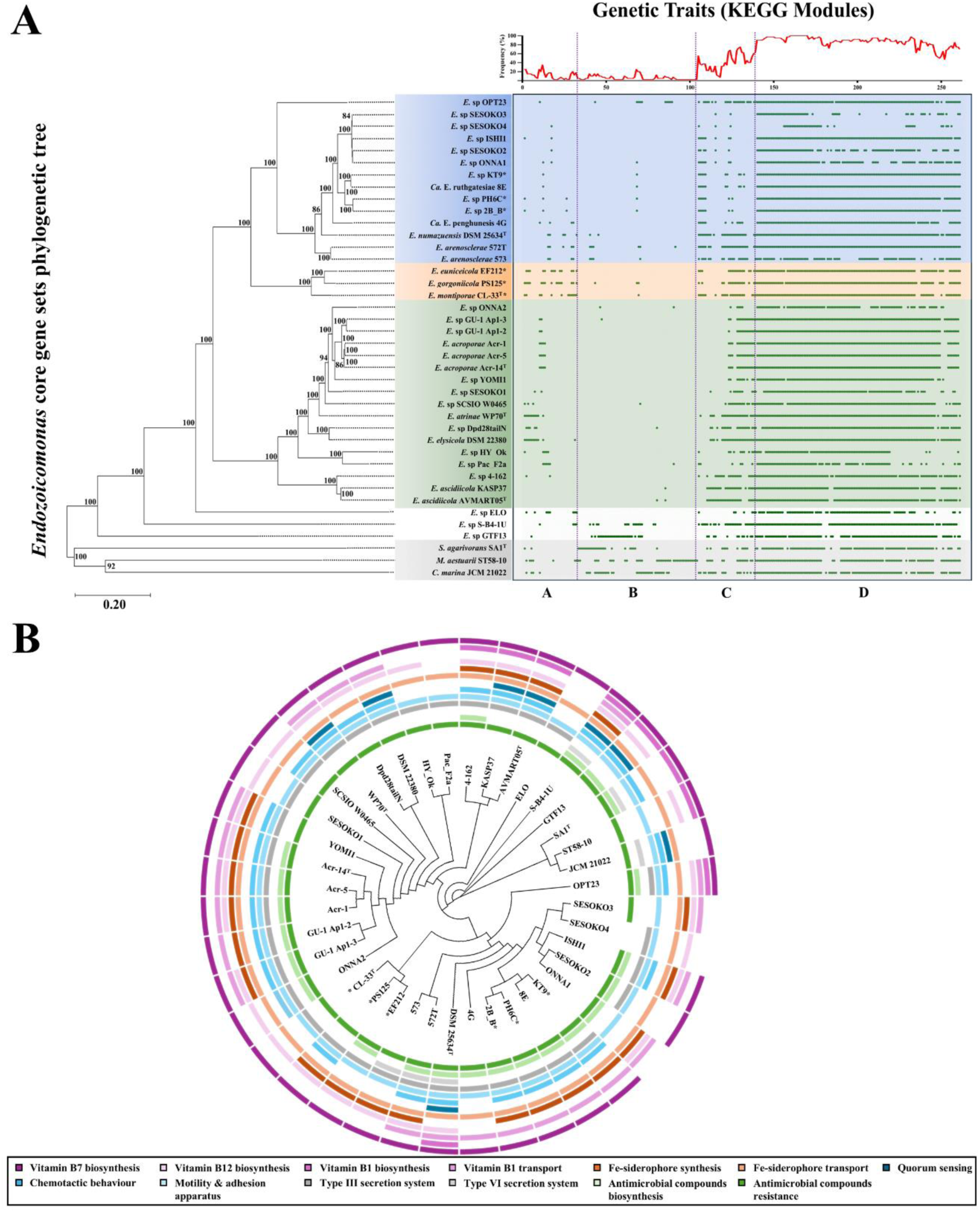
Genetic and interaction traits of 37 *Endozoicomonas* genome assemblies, accompanied by a single-copy core gene phylogenetic tree. (A) *Endozoicomonas* genetic trait frequency constructed with KEGG modules. Blue, orange, and green represent 3 major *Endozoicomonas* clades. White represents known distantly related *Endozoicomonas*, and gray represents non-*Endozoicomonas* outgroups. (B) Radial plot of *Endozoicomonas* interaction traits. Colors represent specific interaction traits in Endozoicomonas. Trait information was extracted from KEGG modules. * indicates *Endozoicomonas* strains sequenced in this study.

To explore potential functional implications of these genetic differences, we analyzed five traits likely involved in microbial and host interactions: vitamin biosynthesis and transport, siderophore synthesis and transport, bacterial communication, secretion systems, and antimicrobial compound biosynthesis and resistance (Supplementary Table S2). A radial plot of interaction traits (Figure 3B) revealed several notable patterns. Most *Endozoicomonas* lack both the ability to synthesize and transport vitamin B12 (81.08%), yet they retain the capacity for vitamin B7 biosynthesis (91.89%), vitamin B1 biosynthesis (56.76%) and vitamin B1 transport (75.68%). However, all *Endozoicomonas* strains retain the ability to transport siderophores (100%), and over half (56.76%) also possess the additional capability to synthesize them. Despite the absence of the quorum sensing trait in most strains (81.08%), *Endozoicomonas* displays traits associated with responding to chemical stimuli (67.57%), independent motility, and adherence (100%) to other bacteria or hosts. The secretion system analysis revealed a predominant reliance on Type III secretion systems (89.19%) for host interactions, with only a minority of strains exhibiting a Type VI secretion system (10.81%). Finally, most *Endozoicomonas* strains exhibit resistance to specific antimicrobial compounds (97.3%), and about half of them produce known antimicrobial compounds (56.76%), potentially providing a competitive advantage over other microbial peers.

### Structural and Functional Heterogeneity of Giant Proteins in *Endozoicomonas*

During functional annotation of *Endozoicomonas* genomes, we identified 92 genes exceeding 15 kbp in 27 of 37 *Endozoicomonas* genomes, each encoding a so-called “Giant Protein” (GP). These genes were classified into 3 size categories: small GPs (>15 kbp; 50 genes), medium GPs (>25 kbp; 41 genes), and a single large GP (>65 kbp; 1 gene) (Table 2). To investigate potential structural similarities, we performed AAI analyses on the small and medium GP subsets separately. The resulting AAI heatmaps revealed that both GP classes could be further subdivided into discrete clusters (Figure 4): medium GPs formed 2 major clusters (Figure 4A), whereas small GPs formed 5 clusters (Figure 4B). Notably, both medium and small GPs could not be clustered together, suggesting strong dissimilarity. Moreover, the distribution of these clusters coincided with the bacterial core gene phylogeny (Figure 2), with most GP clusters occurring in Clade 1.

**Figure 4.**
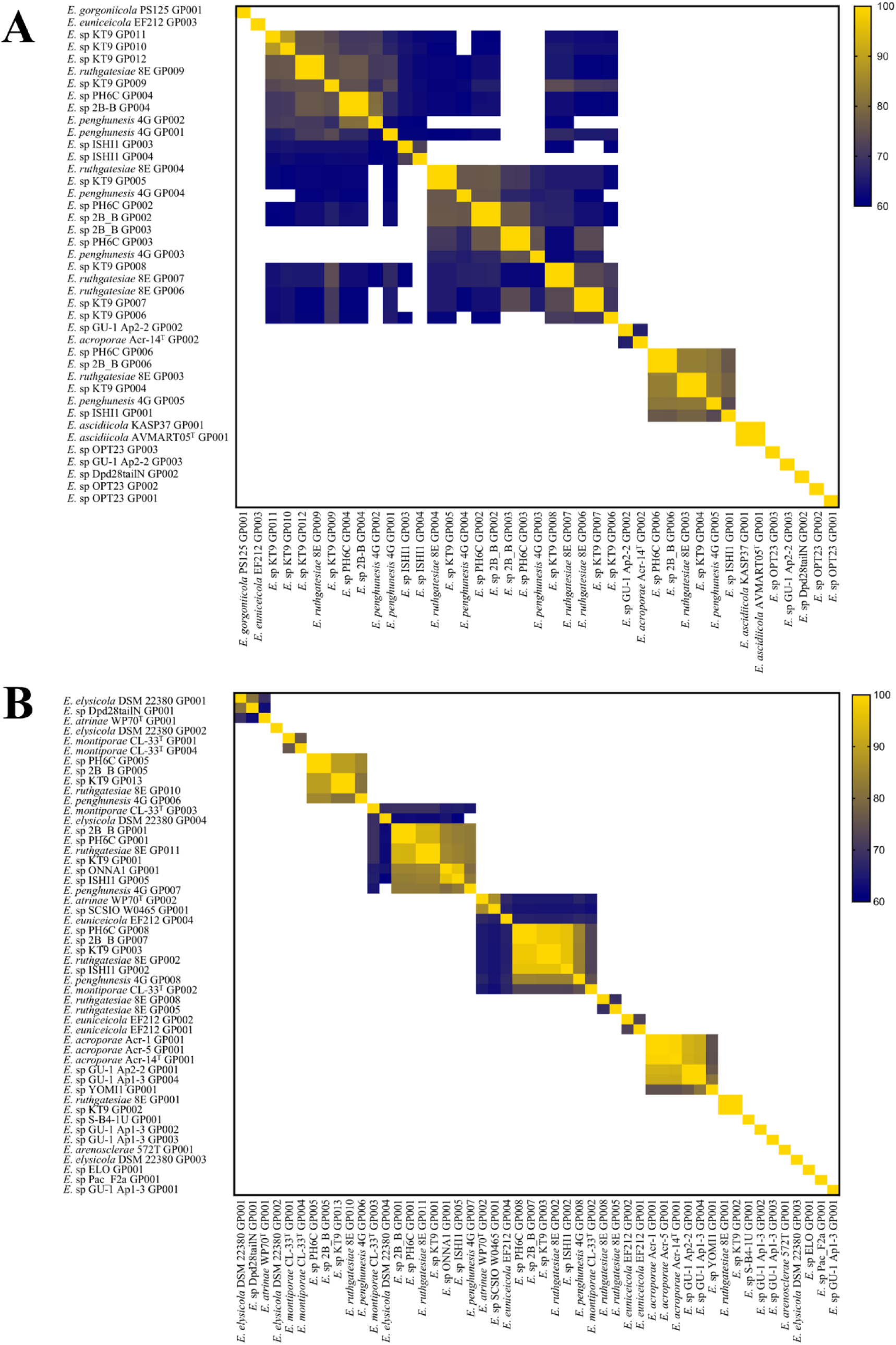
Average Amino Acid Identities (AAIs) of Giant Proteins (GPs). (A) The AAI of medium GPs (>25kbp) giant in *Endozoicomonas* genomes. (B) The AAI of small GPs (> 15 kbp) giant in *Endozoicomonas* genomes. White regions indicate that AAI similarity is < 60%. AAI data were clustered with EzAAI approach.

**Table 2.**
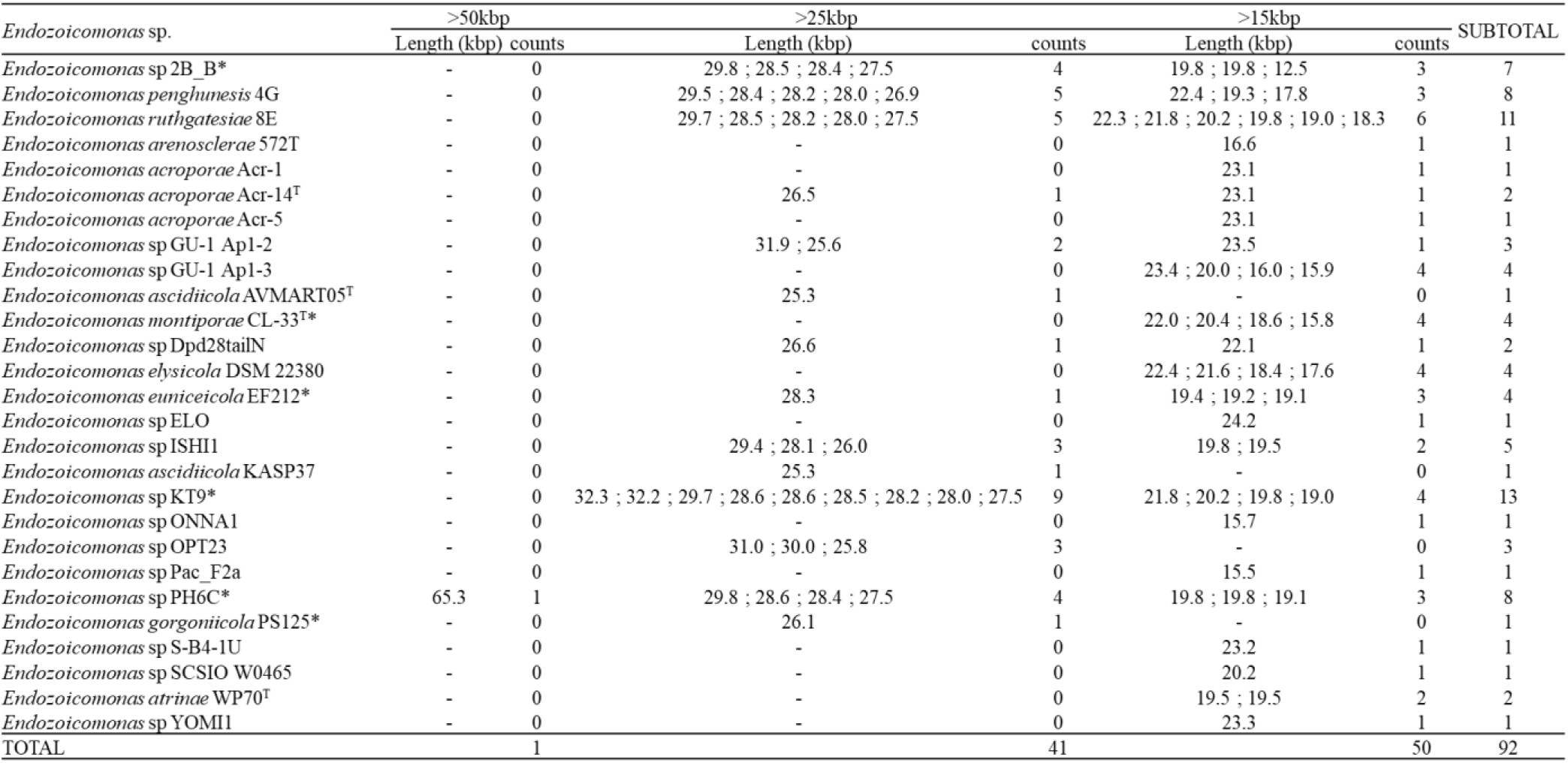
Distribution of various GP classes among 27 *Endozoicomonas* genomes.

To better understand their biological significance, we examined functional domains of each GP class. InterProScan [49] analysis showed that small GP clusters were enriched in domains related to cell adhesion, including HemolysinCabind, VCBS, Laminin_G_3, and Cadherin-related domains (Supplementary Table S3). In contrast, medium GP clusters harbored domains characteristic of exotoxin-associated proteins, such as TcdA/TcdB catalytic glycosyltransferases, CGT/MARTX cysteine protease (CPD) domains, and Peptidase_C80 domains (Supplementary Table S3). The large GP contained multiple non-ribosomal peptide synthetase (NRPS)-related domains, including Phosphopantetheine attachment sites, LCL_NRPS-like, A_NRPS, Cyc_NRPS, AdoMet_Mtases, and McbC_SagB-like_oxidoreductases (Supplementary Table S3). Further antiSMASH [50] analysis also confirmed the presence of NRPS/PKS modules in large GPs that are associated with antimicrobial synthesis. Prediction of large GP 3D structure using AlphaFold3 [51] indicated that NRPS-related domains are surface-exposed, potentially facilitating antimicrobial synthesis (Figure 5). Interestingly, unclustered small and medium GPs exhibited domain compositions similar to those of small and medium GP clusters (Supplementary Table S4), suggesting that they may perform comparable functions despite their amino acid divergence.

**Figure 5.**
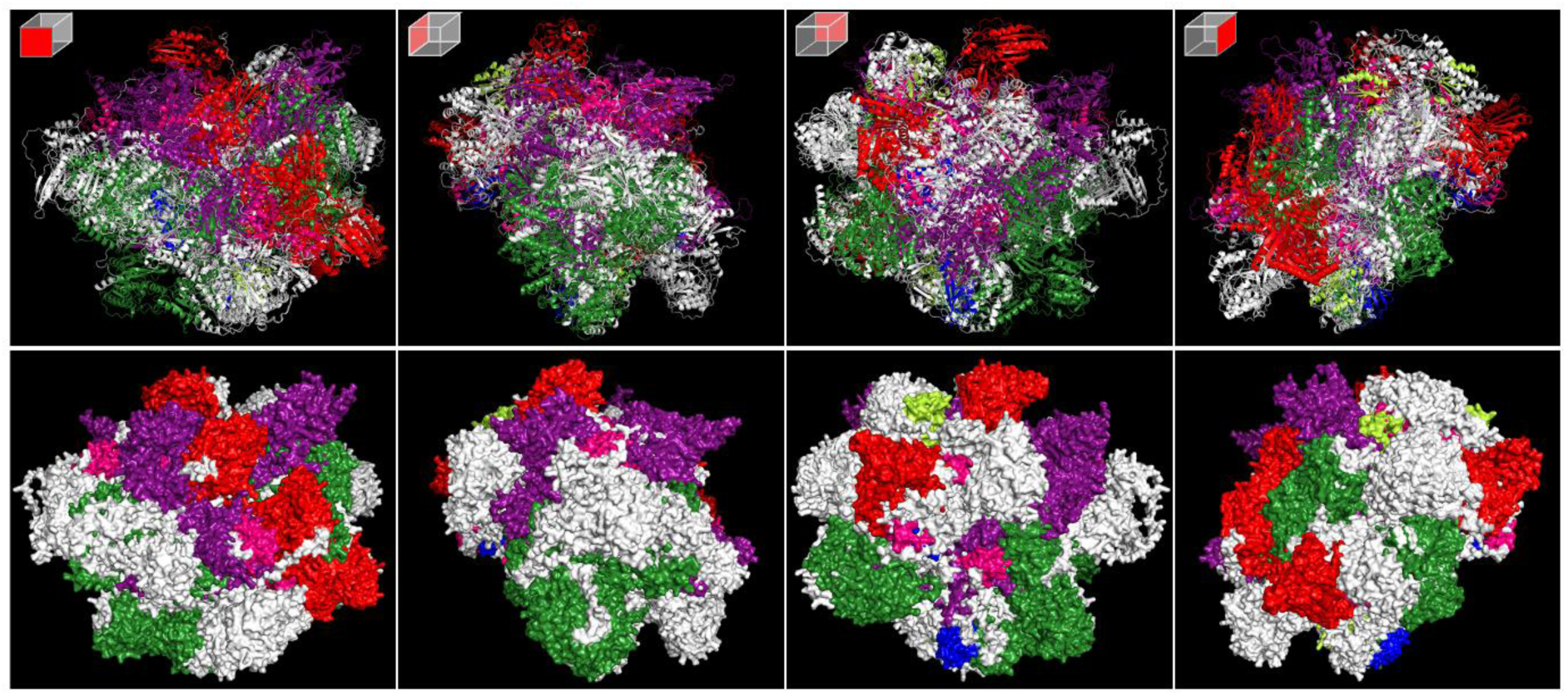
Secondary structure (top) and molecular surface structure (bottom) of a 65-kbp GP in *Endozoicomonas* sp PH6C. Domains predicted by InterProScan are colored differently: Phosphopantetheine attachment site (pink), LCL_NRPRS-like (purple), A_NRPS (green, Cyc_NRPS (red), McbC_SagB-like_oxidoreductase (blue), and AdoMet_Mtases (yellow)

### Diversity and Evolution of Prophages and CRISPR-Cas Systems in *Endozoicomonas*

To explore interactions between prophages and immune defense mechanisms in *Endozoicomonas*, we investigated prophage sequences and CRISPR-Cas systems in 37 genomes. Prophage analysis using PHASTER identified 207 putative prophage regions, categorized as intact, incomplete, or questionable (Supplementary Table S5). Intact prophages were present in 6 newly sequenced *Endozoicomonas* genomes and 16 publicly available *Endozoicomonas* genomes, whereas the other 15 *Endozoicomonas* genomes lacked intact prophages. Strains in Clade 1 contained a single intact prophage, while strains in Clades 2 and 3 harbored multiple intact prophages (Supplementary Table S5). Intact prophages ranged in size from 10 kbp to 75 kbp, significantly larger than incomplete and questionable prophages (Supplementary Figure S3A), and accounted for 0.4%–6.1% of genome content (Supplementary Figure S3B). However, a weak correlation between prophage composition and genome size (R² = 0.38; Supplementary Figure S3C). Analysis of GC content revealed that intact prophages exhibited significantly higher GC percentages than host genomes and other prophage types (Figure 6A). ANI heatmap analysis (Figure 6B) clustered prophages into five major groups, three of which were dominated by Clade 3 strains. Some intact prophages, e.g., from *E.* sp SCSIO W0465, *E.* sp GU1 Ap1-2, and *E. euniceicola* EF212, spanned multiple clusters, indicating acquisition from diverse sources via bacteriophage infection or other gene transfer ways. Phylogenetic analysis using VIPtree suggested that these intact prophages were derived from diverse bacteriophages primarily infecting *Pseudomonadota* (formerly *Proteobacteria*), a major phylum of Gram-negative bacteria (Figure 6C).

**Figure 6.**
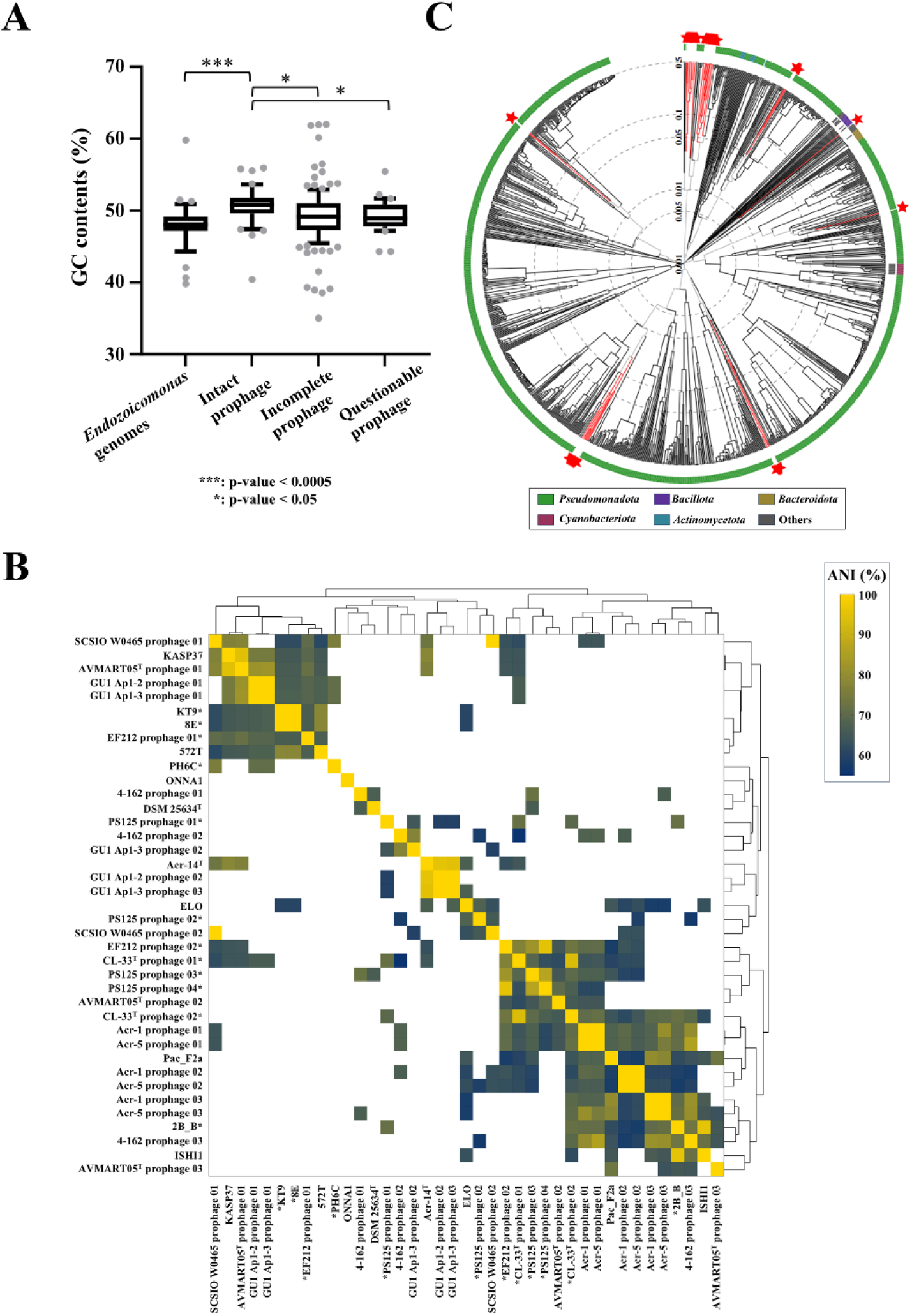
In-depth analysis of intact prophage revealed that *Endozoicomonas* strains acquired intact prophages via HGT. (A) Comparison of GC contents (%) between *Endozoicomonas* and its prophages. (B) ANI heatmap of 40 intact *Endozoicomonas* prophages. ANI data were clustered using a hierarchical clustering approach. The white region indicates that ANI similarity is less than 55%. * indicates *Endozoicomonas* strains sequenced in this study. (C) Phylogeny of *Endozoicomonas* intact prophages associated with other publicly known bacteriophages, constructed by VIPtree. Red stars indicate the intact *Endozoicomonas* prophages identified in this study.

Functional annotation of intact prophage gene content revealed 1,857 genes across 22 *Endozoicomonas* genomes, classified as bacterial-derived genes (76.25%), virus-derived genes (23.59%), and eukaryote-derived genes (0.16%) (Supplementary Figure S4A). COG classification (Supplementary Figure S4B) showed that 57.14% of these genes lacked functional annotation, with the remainder categorized under functions such as replication, recombination, and repair (9.37%, category L); transcription (2.31%, category K); and cell cycle control, division, chromosome partitioning (0.86%, category U). Notably, many unclassified genes (category S, 24.02%) were phage-related proteins.

To complement the prophage analysis, we examined CRISPR-Cas systems of *Endozoicomonas* as they constitute a key defense mechanism against bacteriophage infection. CRISPRCasFinder [56] identified 27 complete CRISPR-Cas systems in 19 *Endozoicomonas* genomes (Supplementary Table S5). The absence of CRISPR-Cas systems in certain genomes could not be attributed to poor assembly quality, as high-quality assemblies such as *Endozoicomonas* sp PH6C also lacked this system. A CRISPR similarity heatmap (Figure 7) revealed 5 distinct CRISPR clusters, which intriguingly showed no correlation with *Endozoicomonas* phylogeny (Figure 1) nor *Endozoicomonas* prophage ANI cluster (Figure 5). Each CRISPR cluster comprised strains from multiple phylogenetic clades, suggesting that CRISPR-Cas system development in *Endozoicomonas* is independent of both prophage acquisition and evolutionary lineage.

**Figure 7.**
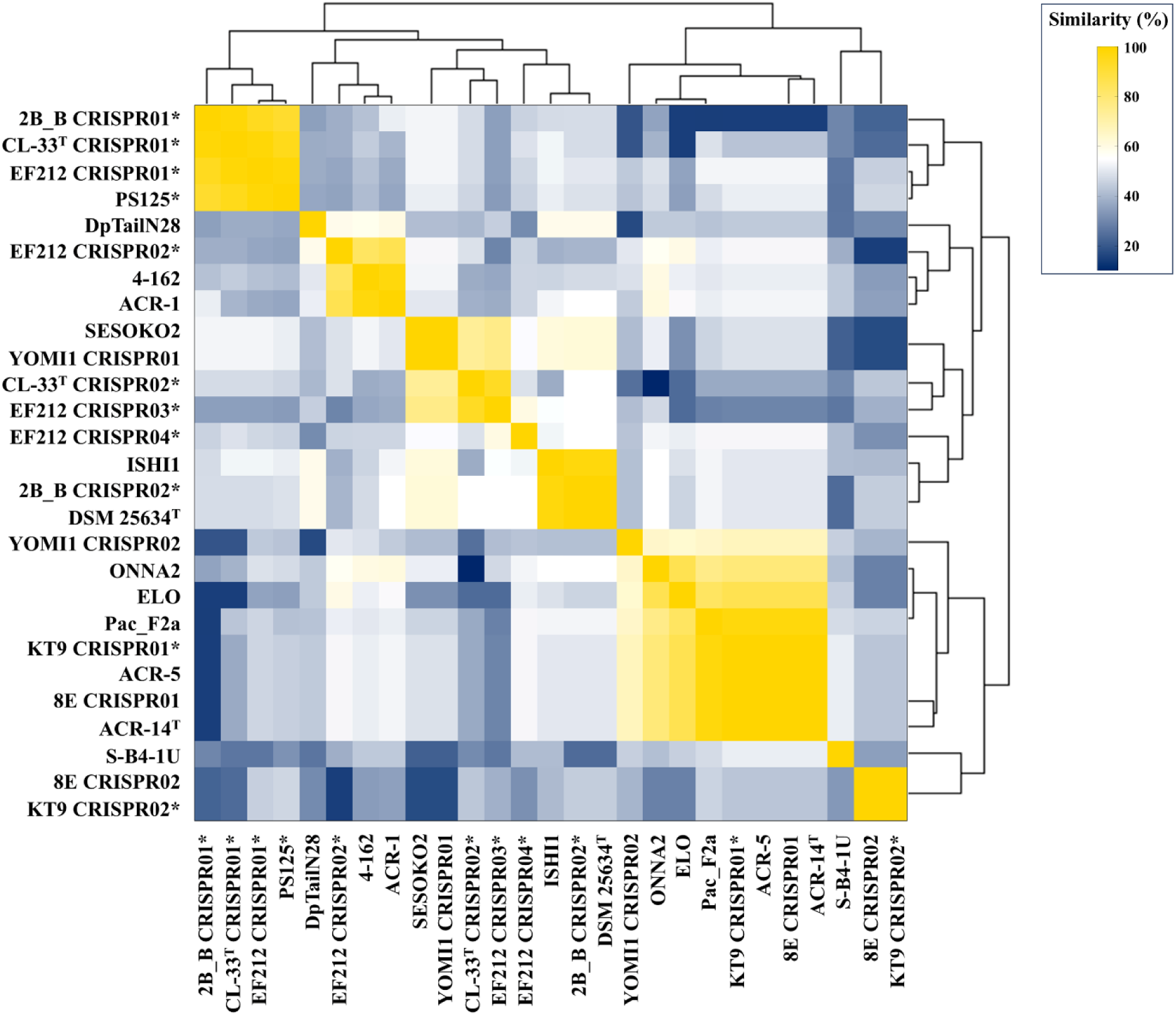
Pair-wise comparison of CRISPR sequences identified in various *Endozoicomonas* genome assemblies. Similarity scores (%) between CRISPR sequences were calculated with emboss. * indicates *Endozoicomonas* strains sequenced in this study.

## Discussion

### High-Resolution Sequencing Unlocks the Genetic and Functional Traits of *Endozoicomonas*

Since the discovery of *Endozoicomonas* in marine slugs [1], this bacterial genus has been identified in numerous marine organisms and linked to diverse functional roles. For instance, *Endozoicomonas* is an integral member of coral holobionts, with many studies underscoring its contribution to coral health and resilience [15, 18, 20, 27]. However, reports of pathogenic or parasitic behavior of certain strains, indicate significant genomic diversity and host-specific adaptations within *Endozoicomonas* [4, 5]. Despite this, efforts to explore these differences have been hindered by limitations of next-generation sequencing technology, which often produces fragmented genome assemblies, restricting pan-genome studies and insights into strain-specific features [10, 26, 27]. In this study, we employed third-generation HiFi sequencing, to overcome these challenges, resulting in complete or nearly complete *Endozoicomonas* genomes. This approach provides an essential framework for *Endozoicomonas* pan-genomic studies, facilitating large-scale comparative analyses to discover gene gain, loss, and adaptation patterns across different strains. More importantly, our high-quality assemblies enable identification of critical genomic features, such as gene clusters, genomic structure and organization for gene regulation, operons, giant proteins, or prophages, which were previously impossible to explore due to fragmented datasets. In this study, we not only sequenced 6 *Endozoicomonas* genomes, but also identified unique genomic features, including previously unreported genetic and interaction traits, giant proteins, and an in-depth investigation of its CRISPR-Cas system and prophages using the most comprehensive pan-genomic analysis conducted to date. These results not only enhance our understanding of *Endozoicomonas,* but also establish new avenues for ecofunctional research.

### Evolutionary Complexity of *Endozoicomonas*: Identification of a Novel Phylogenetic Clade

This comparative analysis and pan-genomic study of 37 *Endozoicomonas* strains revealed remarkable genomic diversity, shedding new light on the evolutionary and functional complexity of this genus. Notably, 3 distinct phylogenetic clades were identified, characterized by unique orthogroup compositions and genetic traits. This finding represents a significant advance, as previous studies had suggested that *Endozoicomonas* phylogeny was confined to 2 clades, with *E. montiporae* CL-33^T^ occupying an intermediate position between them [10, 26–28]. However, our study revealed the emergence of a novel and distinct clade, situated between the previously defined two clades, driven by the inclusion of newly sequenced *E. euniceicola* EF212 and *E. gorgoniicola* PS125, isolated from soft corals. Core gene set analysis and 16S ribosomal phylogeny construction provided robust evidence for this revised phylogenetic framework, suggesting that this new clade may be dominated by *Endozoicomonas* strains that are shared between stony and soft corals, supporting a previous report that identified *Endozoicomonas* as being shared between these coral types in the deep sea [59]. These findings indicate that the diversity of *Endozoicomonas* is much higher than previously thought, suggesting a novel evolutionary trajectory. It also emphasizes the importance of exploring underrepresented strains to uncover hidden diversity in microbial lineages.

### Deciphering Interactions and Functional Roles of *Endozoicomonas* in Holobionts

The ability of *Endozoicomonas* strains to interact with hosts and other microbes depends upon specific genetic traits that enable mutualism [60]. Previous studies have highlighted the functional specificity of *Endozoicomonas*, with different strains contributing distinct nutrients essential for coral health. For instance, comparative genomic analyses provide indirect evidence that *Endozoicomonas* may contribute to chitin degradation [30, 61], cycling of carbohydrates, proteins, and amino acids, and biosynthesis of secondary metabolites such as hormones, cofactors, and vitamins that support hosts or other holobiont members [10, 27, 28, 62, 63]. Additionally, experimental evidence indicates that *E. acroporae* and *E. ruthgatesiae* 8E metabolize dimethylsulfoniopropionate (DMSP) [23, 29], whereas *E. montiporae* CL-33^T^ is associated with steroid degradation [22]. Our pan-genomic study revealed that the 3 *Endozoicomonas* clades exhibit clear genetic trait specificity (Figure 3A), with certain traits linked to specific clades. However, a more complex and nuanced picture emerged when examining interaction traits of *Endozoicomonas* (Figure 3B), shedding light on how these bacteria interact with their hosts and surrounding microbes (Figure 8).

**Figure 8.**
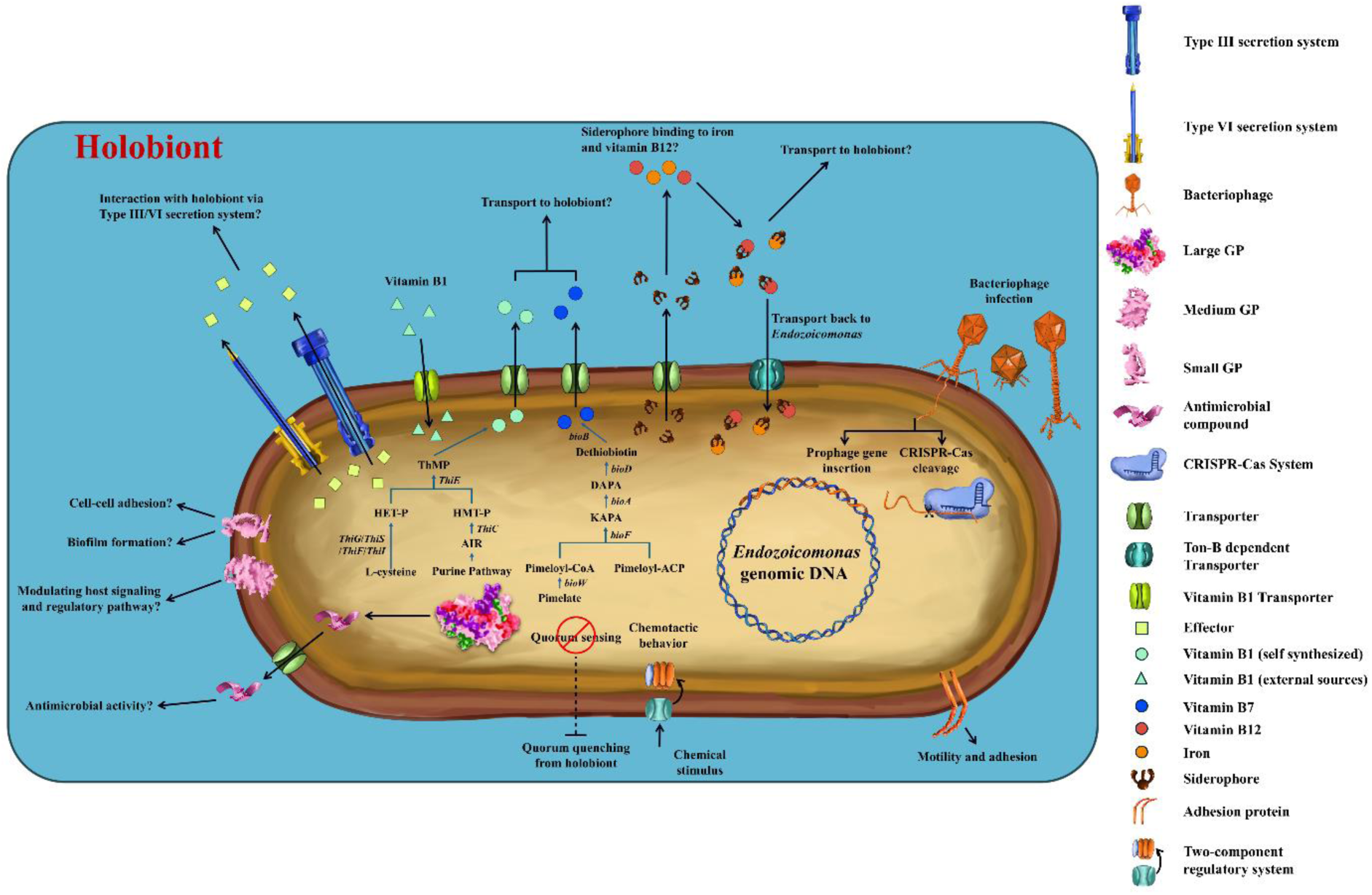
Genomic features of *Endozoicomonas* strains involved in interactions with hosts and surrounding microbes. This schematic representation highlights key genetic traits, including giant proteins, prophages, and the CRISPR-Cas system, and illustrates how these features influence host-microbe and microbe-microbe interactions. Abbreviations of chemical compounds in vitamin B1 and B7 biosynthesis are: AIR, 5-aminoimidazole ribotide; HMP-P, 4-amino-5-hydroxymethyl-2-methylpyrimidine phosphate; HET-P, 4-Methyl-5-Hydroxyethylthiazole Phosphate; ThMP, thiamine monophosphate; KAPA, 7-Keto-8-Aminopelargonate; DAPA, 7,8-diaminopelargonic acid

Interaction of *Endozoicomonas* strains with hosts and other microbes revealed that they retain the ability to synthesize vitamin B1 (thiamine), which is consistent with previous *Endozoicomonas* proteome observations [28]. Moreover, *Endozoicomonas* also retains the capacity to synthesize vitamin B7, a cofactor important for marine bacterial metabolism and growth [64]. However, *Endozoicomonas* lacks the capacity to produce or transport vitamin B12 (cobalamin) (Figure 8). Vitamin B12 is a critical cofactor for DNA synthesis and fatty acid and amino acid metabolism in both eukaryotes and prokaryotes [65]. Since most marine hosts cannot synthesize vitamin B12, they rely on microbial symbionts [66]. The absence of vitamin B12 biosynthesis in *Endozoicomonas* demonstrates that these bacterial symbionts do not contribute this essential nutrient. Similar deficiencies in vitamin B12 production have been observed in other endosymbiotic bacteria, such as *Buchnera aphidicola* [67] and *Wigglesworthia glossinidia* [68], though these examples are restricted to insect hosts. The absence of vitamin B12 production in *Endozoicomonas* raises intriguing questions about how this important bacterial group acquires this essential nutrient, and what interactive relationships exist between *Endozoicomonas* and other holobiont members.

Our findings indicate that *Endozoicomonas* may provide alternative nutritional or functional benefits to hosts. A key observation was that half of *Endozoicomonas* strains able to synthesize siderophores, underscoring their adaptability to nutrient-limited marine environments. Siderophores facilitate iron acquisition, a process critical for cellular functions yet challenging in the often iron-scarce oceanic ecosystem [69]. This trait is particularly advantageous in oligotrophic waters, where efficient iron acquisition confers a competitive edge [70]. Previous studies have demonstrated the ecological significance of siderophore production in marine bacteria. For instance, siderophore production by *Alteromonas macleodii* enhances iron availability for co-existing marine organisms, including phytoplankton, thereby supporting marine ecosystems [71]. Similarly, *Endozoicomonas* could thrive in oligotrophic waters and supply iron to hosts or other microbes.

Notably, all *Endozoicomonas* strains possess the TonB-dependent receptor system, including TonB, ExbB, and ExbD, which is critical for siderophore transport. This system may serve as an alternative mechanism for scavenging vitamin B12 from the surrounding environment, compensating for the absence of a dedicated vitamin B12 transport protein, as in most *Endozoicomonas* strains [72–75]. This dual functionality of siderophores, facilitating both iron acquisition and potentially vitamin B12 scavenging, underscores the ecological versatility of *Endozoicomonas*.

Quorum sensing is a crucial bacterial cell-cell communication process that enables bacteria to regulate gene expression and biofilm formation in response to population density [76]. Many *Endozoicomonas* strains lack this trait, suggesting that they rely on alternative, unidentified mechanisms for bacterial communication. This absence of quorum sensing also implies that neither hosts nor microbes employ quorum quenching, a strategy that disrupts or inhibits bacterial quorum sensing [77, 78] against *Endozoicomonas* strains. The deficiency of quorum sensing may provide a selective advantage, allowing *Endozoicomonas* to evade disruption or manipulation by hosts and other microbes. However, further investigations and experimental validation are necessary to confirm this hypothesis.

### Functions of Small, Medium, and Large Giant Proteins in *Endozoicomonas* Biology

Genes encoding giant proteins (GPs) ≥ 5 kbp are rare in bacteria, constituting less than 0.2% of bacterial genomes due to their high energy costs for synthesis [79]. The presence of such genes suggests that they may fulfill critical, yet underexplored, biological roles. Most of these genes encode GPs associated with surface proteins involved in cell attachment or with non-ribosomal peptide synthetases (NRPSs) and polyketide synthases (PKSs), which are responsible for synthesizing non-ribosomal peptides and polyketides [79]. These compounds have demonstrated diverse bioactivities, including antimicrobial, antifungal, antiparasitic, antitumor, and immunosuppressive properties. In our study, the large GP identified in *Endozoicomonas* sp. PH6C aligns with these functional profiles. AntiSMASH analysis revealed several NRPS regions in the large GP, suggesting potential inhibition of gram-positive bacteria [80] and biosurfactant activity [81]. However, as only a single large GP was identified in *E.* sp. PH6C, we hypothesize that this unique GP may possess antimicrobial properties, allowing *Endozoicomonas* to compete with other microbes.

Interestingly, our findings also revealed that the majority of *Endozoicomonas* strains in Clades 1 and 2 are enriched in a variety of clustered small and medium GPs, suggesting that these genes are essential to the genus. Further analysis showed that clustered small and medium GPs possess distinct domain repeats, likely contributing to diverse biological functions (Figure 8). For instance, these clustered small GPs containing VCBS domain repeats may facilitate cell-cell adhesion [82] or biofilm formation [83], while HemolysinCabind and Laminin_G_3 domains are associated with calcium-binding, which impacts protein structural stability [84, 85]. These findings support the hypothesis that small GPs could contribute to biofilm formation and cell-cell adhesion. Clustered Medium GPs, on the other hand, contain exotoxin-related domains, such as TcdA/TcdB catalytic glycosyltransferases and CGT/MARTX cysteine protease (CPD) domains. These domains typically damage host cells or disrupt cellular metabolism [86]. However, the absence of combined repetitive oligopeptides (CROPs), which are necessary for receptor binding and initiating virulence effects, suggests that these exotoxins are non-functional in clustered medium GPs and are therefore non-toxic to hosts. We speculate that these medium GPs may instead modulate host signalling and regulatory pathways, facilitating *Endozoicomonas* entry into hosts. Meanwhile, unclustered small and medium GPs contain various domains similar to those in the large, clustered medium, and clustered small GP groups, suggesting that these unclustered GPs perform diverse functions despite their amino acid divergence.

### Deciphering Phage-Host Dynamics: Prophage Integration and CRISPR-Cas Interactions in *Endozoicomonas* Genomes

The dynamic interplay between bacteria and bacteriophages in marine environments is a cornerstone of oceanic ecosystems. Bacteriophages, as bacterial predators, employ two replication strategies: lytic and lysogenic [87, 88]. During the lytic cycle, phages infect bacterial hosts, replicate inside them, and ultimately cause cell lysis, releasing a wealth of organic matter and nutrients back into the marine environment. This process contributes to the microbial loop, a critical mechanism that redistributes carbon and nutrients, thereby influencing biogeochemical cycles of the ocean [89, 90]. In contrast, the lysogenic cycle allows bacteriophages to integrate their genetic material into bacterial genomes as prophages via infections or other gene transfer methods [91]. This integration fosters genetic diversification, potentially enhancing bacterial adaptability to fluctuating oceanic conditions [92]. To counter phage predation, bacteria have evolved a sophisticated adaptive immune system known as the CRISPR-Cas system [93]. This mechanism incorporates short DNA sequences from invading phages into bacterial genomes as spacers in CRISPR arrays. These spacers serve as a genetic memory, enabling the bacterial CRISPR-Cas machinery to recognize and cleave invading phage DNA upon subsequent attacks [94].

In this study, we investigated interactions between prophages and the CRISPR-Cas system in *Endozoicomonas* genomes, uncovering intriguing insights. Prophages in *Endozoicomonas* exhibit a significantly higher GC content than their host genomes, suggesting acquisition via bacteriophage infection or other gene transfer methods. Phylogeny indicates that these prophages originate from distant bacteriophage lineages that primarily infect *Pseudomonadota* (*Proteobacteria*). Prophage ANI reveals that these prophages form unique clusters independent of the *Endozoicomonas* core gene set or 16S phylogeny, indicating individual *Endozoicomonas* lineages had different infection histories due to specific hosts or environments (including hosts). These findings point to the possibility that *Endozoicomonas* prophages may employ the “Piggyback-the-Winner” strategy, in which the lysogenic bacteriophage adopts a temperate lifecycle to avoid predation pressure while integrating into bacterial genomes [95, 96]. This integration not only confers superinfection exclusion to bacteria, but also highlights the strategic localization of lysogenic prophages in a eukaryotic host. For example, Silveira and Rohwer (2016) hypothesized that bacteriophage infection or other gene transfer activities leading to prophage acquisition in *Endozoicomonas* strains, predominantly occurred in the coral’s mucus layer, an environment rich in microbial communities [95]. Notably, the weak correlation between prophage composition and *Endozoicomonas* genome size (Supplementary Figure S3C) strongly suggests that prophage acquisition is not a major driver of genome expansion in *Endozoicomonas*.

Additionally, our findings indicate that the CRISPR-Cas system in *Endozoicomonas* operates independently of prophage acquisition. The extensive diversity of CRISPR sequences across *Endozoicomonas* genomes shows no correlation with the diversity of prophage sequences, suggesting that *Endozoicomonas* acquires CRISPR spacers primarily from lytic rather than lysogenic phages. This is consistent with previous studies, highlighting the selective pressure exerted by lytic phages on the CRISPR-Cas machinery [97, 98]. These findings reveal a complex phage-bacteria network that may influence marine host health and surrounding microbial communities, as proposed by several studies [99–102]. Our result reveals that if *Endozoicomonas* strains do not undergo lysis during bacteriophage infection, injected bacteriophage DNA could face two fates: integration into the *Endozoicomonas* genome as prophage or cleavage by the CRISPR-Cas system (Figure 8). This hypothesis suggests that bacteriophages may help to shape the genomic diversity and adaptability of *Endozoicomonas* in marine hosts, but further investigation and experimental validation are needed.

## Conclusions

In summary, this study presents novel *Endozoicomonas* genome assemblies and also conducts the most comprehensive *Endozoicomonas* pan-genomic analysis to date, uncovering previously unknown genomic features. Our findings suggest that *Endozoicomonas* strains contribute alternative nutrients to support holobionts, compensating for their inability to synthesize vitamin B12. Additionally, the absence of quorum-sensing in *Endozoicomonas* implies the use of an alternative, unidentified mechanism for bacterial communication or biofilm formation, potentially enabling these bacteria to evade quorum-quenching inhibition. Furthermore, the discovery of enriched giant proteins in *Endozoicomonas* suggests that these proteins may participate in cell-cell adhesion, biofilm formation, host signalling modulation, and antimicrobial activity. Lastly, in-depth analyses of prophages and the CRISPR-Cas system reveal a complex phage-bacteria network that may influence holobiont health and development. This study provides a valuable foundation for future investigations into ecofunctional traits of *Endozoicomonas* and their broader ecological significance.

## List of abbreviations

GPs: Giant Proteins

## Declarations

### Ethics approval and consent to participate

Not applicable.

### Consent for publication

Not applicable.

### Data availability

The six *Endozoicomonas* genome assemblies and raw read data used in this study have been deposited at the National Center for Biotechnology Information (NCBI) BioSample under the BioProject PRJNA1230818 (https://www.ncbi.nlm.nih.gov/bioproject/PRJNA1230818) and PRJNA871056 (https://www.ncbi.nlm.nih.gov/bioproject/PRJNA871056).

### Author’s contributions

SL Tang designed and supervised the project. SL Lim wrote the manuscript. SL Lim and CH Chin performed the bioinformatic analysis. YJ Chiou, MT Hsu, PW Chiang, HJ Chen and YC Tu contributed to the sample collection, isolation and sequencing experiments. The authors read and approved the final manuscript.

## Acknowledgements

We thanked Academia Sinica, Taiwan, for funding the research project and PacBio sequencing.

---

**Fig. S1.**
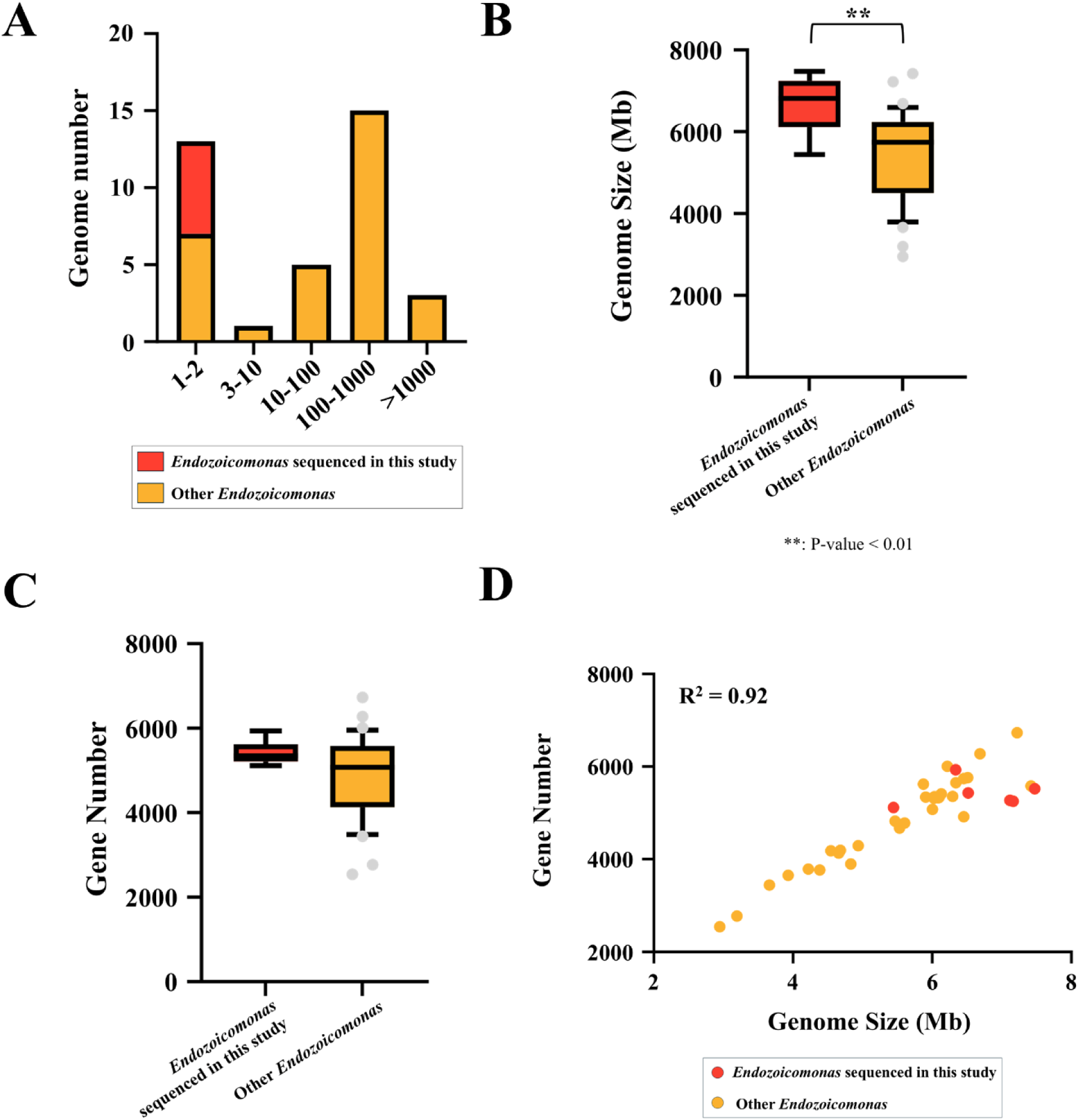
Comparative analysis of 6 *Endozoicomonas* genomes sequenced in this study with 31 publicly available *Endozoicomonas* genome assemblies. **(A) Contig numbers in various *Endozoicomonas* genome assemblies.** The 6 sequenced *Endozoicomonas* genome assemblies have the lowest contig numbers, indicating high-quality assemblies, whereas other publicly available *Endozoicomonas* genome assemblies varied from 1 or 2 to more than 1000 contigs. (B) **Genome sizes of 6 sequenced *Endozoicomonas* genomes and other publicly available *Endozoicomonas* genome assemblies**. Student’s two-tailed t-test revealed significant differences between the 31 publicly available *Endozoicomonas* genome sizes and those sequenced in this study. **(C) Gene content of 6 sequenced *Endozoicomonas* genomes and other publicly available *Endozoicomonas* genome assemblies**. *Endozoicomonas* nucleotide sequences were annotated using Prokka. **(D) Correlation between** *Endozoicomonas* genome sizes and gene numbers. Correlation analysis was performed with Pearson’s correlation.

**Fig. S2.**
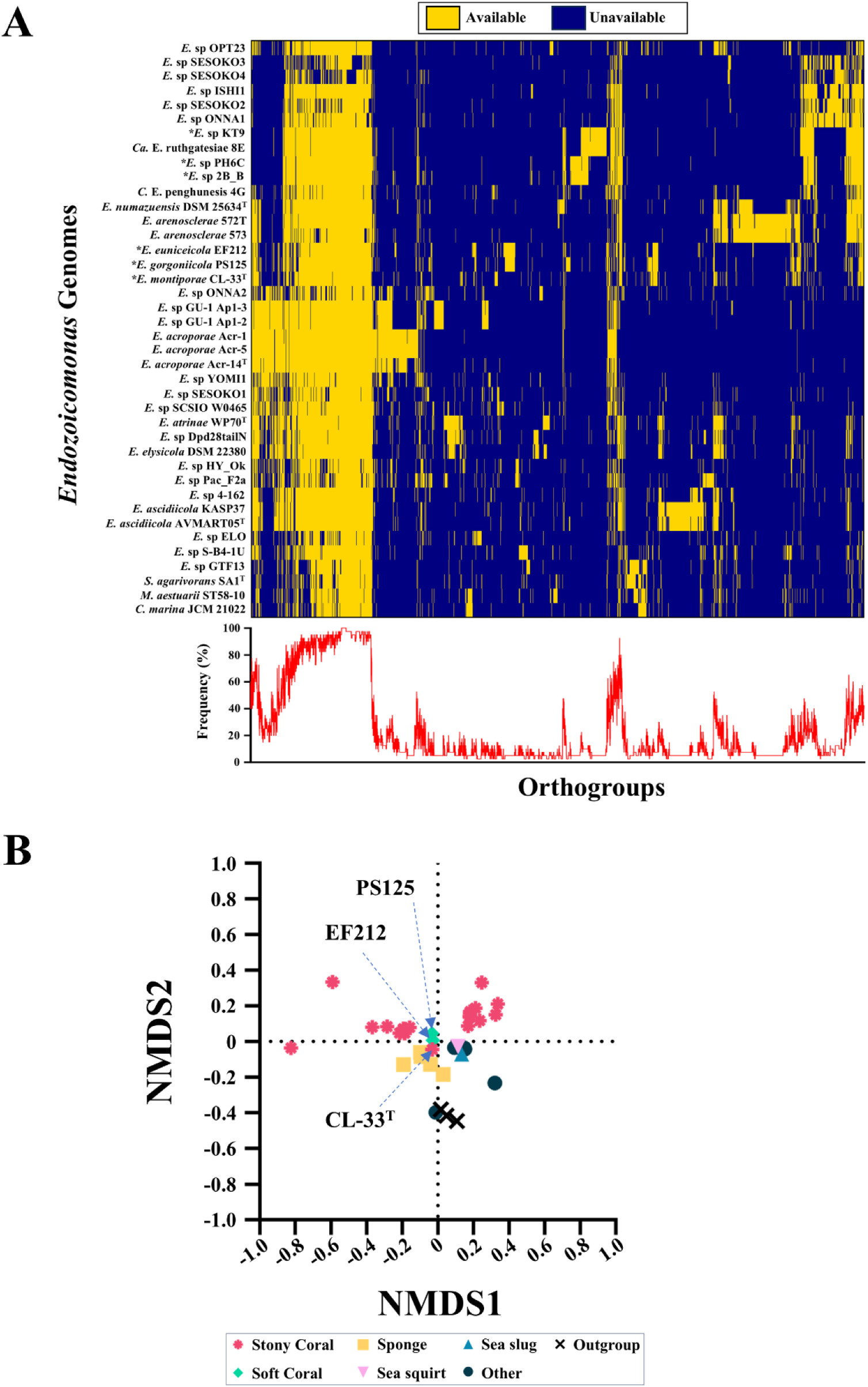
Pan-genomic analysis of 37 *Endozoicomonas* genome assemblies. **(A) The 37 *Endozoicomonas* orthogroup heatmap.** Hierarchical clustering of orthogroups was based on the single-copy core gene phylogenetic tree, and each orthogroup’s appearance frequency (%) on *Endozoicomonas* was calculated. (B) **Non-metric Multi-dimensional Scaling (NMDS) plot of 37 *Endozoicomonas* genome assemblies.** NMDS was calculated using the R vegan package, and the stress value was 0.11, indicating the NMDS analysis is good fit.

**Fig. S3.**
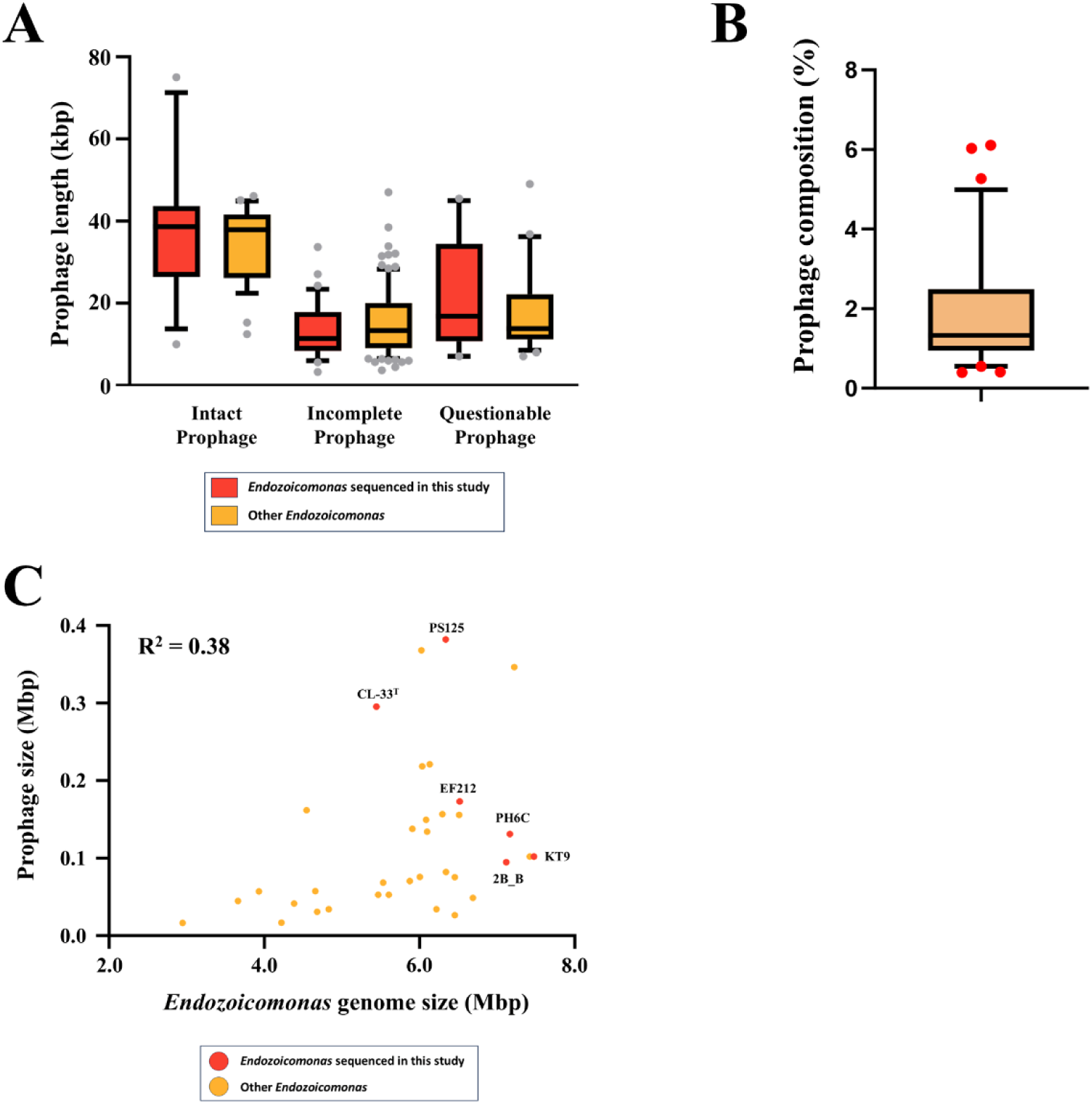
Statistical analysis of intact, incomplete, and questionable prophages identified in 37 *Endozoicomonas* genome assembles. **(A) Lengths of prophages (kbp) identified in *Endozoicomonas* genomes**. The red box plot presents the 6 *Endozoicomonas* genomes sequenced in this study. The orange box represents the 31 publicly available *Endozoicomonas* genome assembles. **(B) Prophage composition (%) in 37 *Endozoicomonas* genome assemblies**. Prophage composition includes intact, incomplete, and questionable prophages identified with PHASTER. **(C) Correlation of *Endozoicomonas* genome size (Mbp) and Prophage size (Mbp)**. Correlation analysis was performed with Pearson’s correlation.

**Fig. S4.**
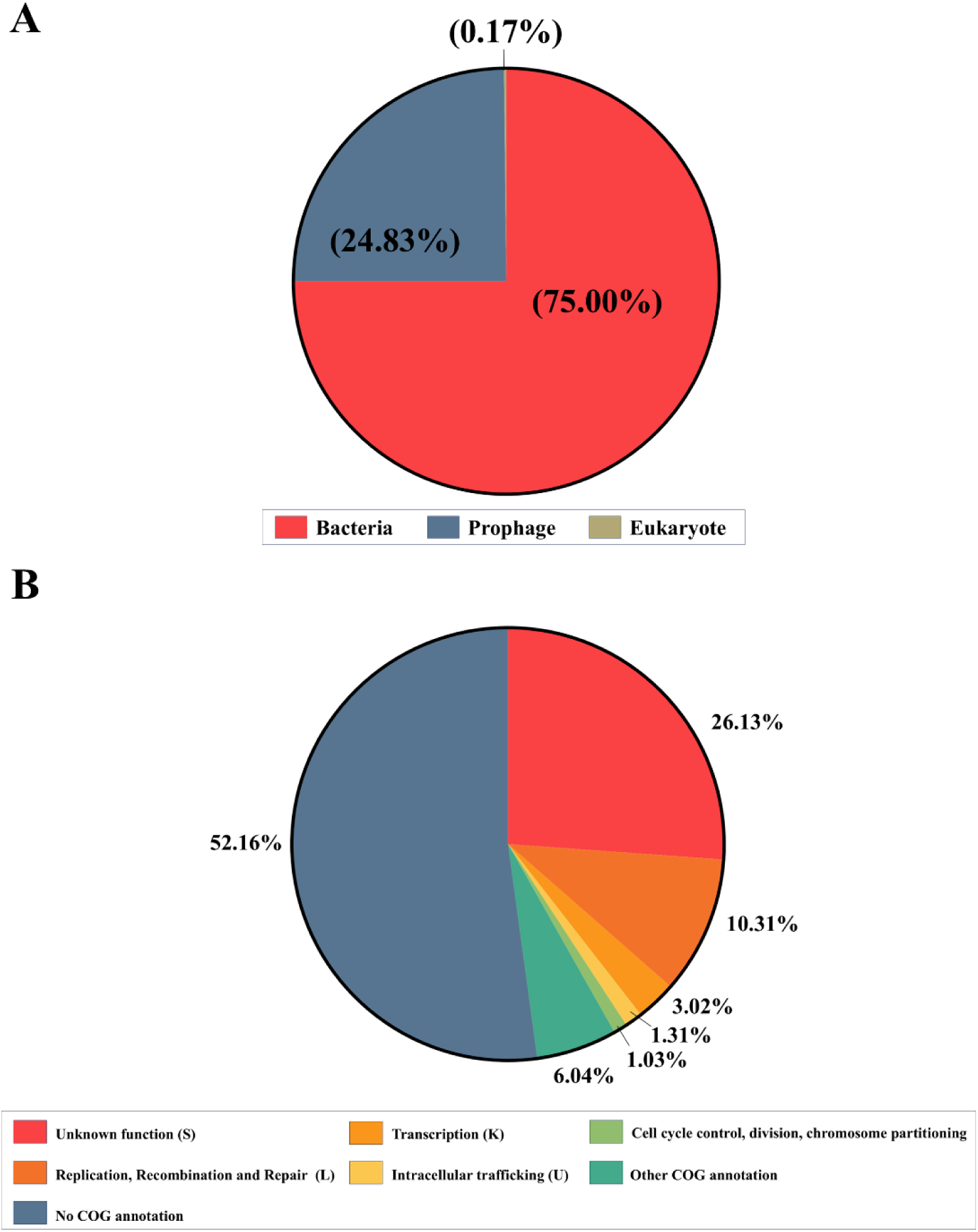
Identification and functional classification of intact *Endozoicomonas* prophage genes. **(A) Origin of gene contents in intact *Endozoicomonas* prophages**. Gene sequences and origins (bacteria, prophage, eukaryote) are classified with BLASTP **(B) COG functional classification of intact *Endozoicomonas* genes**. The COG category classification is performed using eggNOG-mapper.

